# Concurrent action observation but not motor imagery modulates interhemispheric inhibition during physical execution

**DOI:** 10.1101/2023.12.19.572434

**Authors:** Kyle A. Vallido, Matthew W. Scott, Carrie M. Peters, Kelly Spriggs, Nicola J. Hodges, Sarah N. Kraeutner

**Author notes:** **Address for Correspondence** Dr. S.N. Kraeutner, Department of Psychology, the University of British Columbia Okanagan Campus, Rm 204, Arts and Sciences (ASC) 1147 Research Rd, Kelowna, BC Canada V1V 1V7.

## Abstract

Motor control relies on an inhibitory connection between the motor cortices of the brain, known as interhemispheric inhibition (IHI). This phenomenon is well established during the execution of unilateral motor tasks. It is unknown if the neurophysiological effects associated with IHI during physical execution (PE) also occur during action observation (AO) and motor imagery (MI) and/or if the addition of these covert processes to PE moderates IHI; speaking to differences in neurophysiology and functional equivalence. Participants (N=23) performed unilateral concentric wrist contractions (50% maximum voluntary contraction) under three conditions: PE alone, concurrent PE+AO, and concurrent PE+MI. To index IHI, we induced an ipsilateral silent period (iSP) and assessed iSP duration during each condition via neuro-navigated single-pulse transcranial magnetic stimulation (TMS) over the ipsilateral motor cortex. Relative to PE alone, iSP decreased during PE+AO, yet only when this condition preceded PE+MI. iSP duration was not modulated during PE+MI. Together, these data suggest that PE+AO promotes bilateral recruitment and ‘interhemispheric cooperation’ rather than inhibition. AO and MI differentially impact interhemispheric coordination, serving to suppress inhibition only when AO is primed by MI.

---

Covert actions, including action observation (AO; watching the performance of a movement) and motor imagery (MI; the mental rehearsal of a movement), have been argued to be “functionally equivalent” to overt action (Jeannerod, 1994; 2001) and thus rely on an internal simulation of the same action representation as that which governs physical execution. Evidence in support of this “equivalence” stems primarily from neuroimaging work, showing overlap in the neural processes recruited during covert and overt action (see Caspers et al., 2010; Di Rienzo et al., 2016; Hardwick *et al*., 2018) and a wealth of behavioural evidence showing that covert action, observational and/or imagery practice, can effectively drive motor learning (for reviews see Buchignani et al., 2019; Di Rienzo et al., 2016; Eaves et al., 2016; Hodges 2017; Ladda et al., 2021; Ramsey et al., 2021). Despite the emphasis on equivalence between covert and overt action states (Vogt et al., 2013), neural differences have been highlighted in comparative work. Moreover, in a recent theory of covert actions, it has been suggested that there is only equivalence during the planning but not the “execution” component of covert and overt movements, with the former showing a strong reliance on cognitive processes not evidenced during actual action (e.g., Glover & Baran, 2017; Glover et al., 2020; Frank et al., 2023).

With respect to patterns of brain activation, there is work showing that MI leads to activation that is distributed across a number of frontoparietal regions in the brain, with learning through MI leading to changes in connectivity localized to a distributed frontoparietal network (rather than focal changes within a cortical-cerebellar network as in physical execution; Kraeutner et al. 2022). Further, in contrast to physical execution, inconsistencies in contralateral motor cortex activation are reported across studies investigating unilateral movements performed during MI (see Hétu et al., 2013 for a review). During AO of unilateral movements, bilateral recruitment of the motor cortices has been observed (Hardwick et al., 2018), potentially associated with external stimuli processing (Alhaijri et al., 2018; Caspers et al., 2010). Evidence regarding similarities (and differences) between these various action states in terms of cross-hemisphere control has to date been inferred from functional magnetic resonance imaging or positron emission tomography measures (Caspers et al., 2010; Hardwick et al., 2018; Hétu et al. 2013). Yet, while rich spatial information is provided, given these modalities rely on indirect measures of brain activity (whereby neuronal function is inferred rather than directly tested) with low temporal resolution (Cumming et al., 2014; Sutton et al., 2009) we are unable to disentangle execution related processes from those related to planning, with only the latter thought to be based on a shared action representation (Glover & Baran, 2017). To date, there has been no direct testing of cross-hemispheric patterns of cortical control during physical execution that is time-locked to the movement and paired with AO and MI, which could speak better to the equivalence of mechanisms underpinning covert action states.

Physical execution of a unilateral movement relies on inhibitory communication between the two motor cortices in the brain, termed interhemispheric inhibition (IHI; Perez & Cohen, 2009). An important component of single limb actions is the inhibition of one hemisphere in order to effectively monitor/control movements associated with the other hemisphere and relevant effector (e.g., Fling & Seidler, 2012). One way to index IHI is to measure the latency of the ipsilateral silent period (iSP), which is the onset and offset of the inhibitory signal when performing a unilateral contraction (Kuo et al., 2017; Hupfeld et al., 2020). As illustrated in Figure 1a, IHI is assessed using transcranial magnetic stimulation (TMS) delivering a single-pulse over the ipsilateral motor cortex during an active contraction (i.e., rather than over the motor cortex contralateral to the effector). A suprathreshold TMS pulse delivered over the ipsilateral motor cortex results in a brief suppression of GABAergic interneurons supporting the unilateral contraction, reflected by a temporary rapid ‘break’ in the electromyographic (EMG) signal of the contracted muscle. Increasing IHI (indexed by increased iSP duration obtained from the EMG trace) during unilateral movement, has been suggested to reflect sustained lateralized cortical activity. As such, there is reduced interference between the left and right motor cortices which is thought to facilitate precise contralateral motor control (Fling & Seidler, 2012).

**Figure 1.**
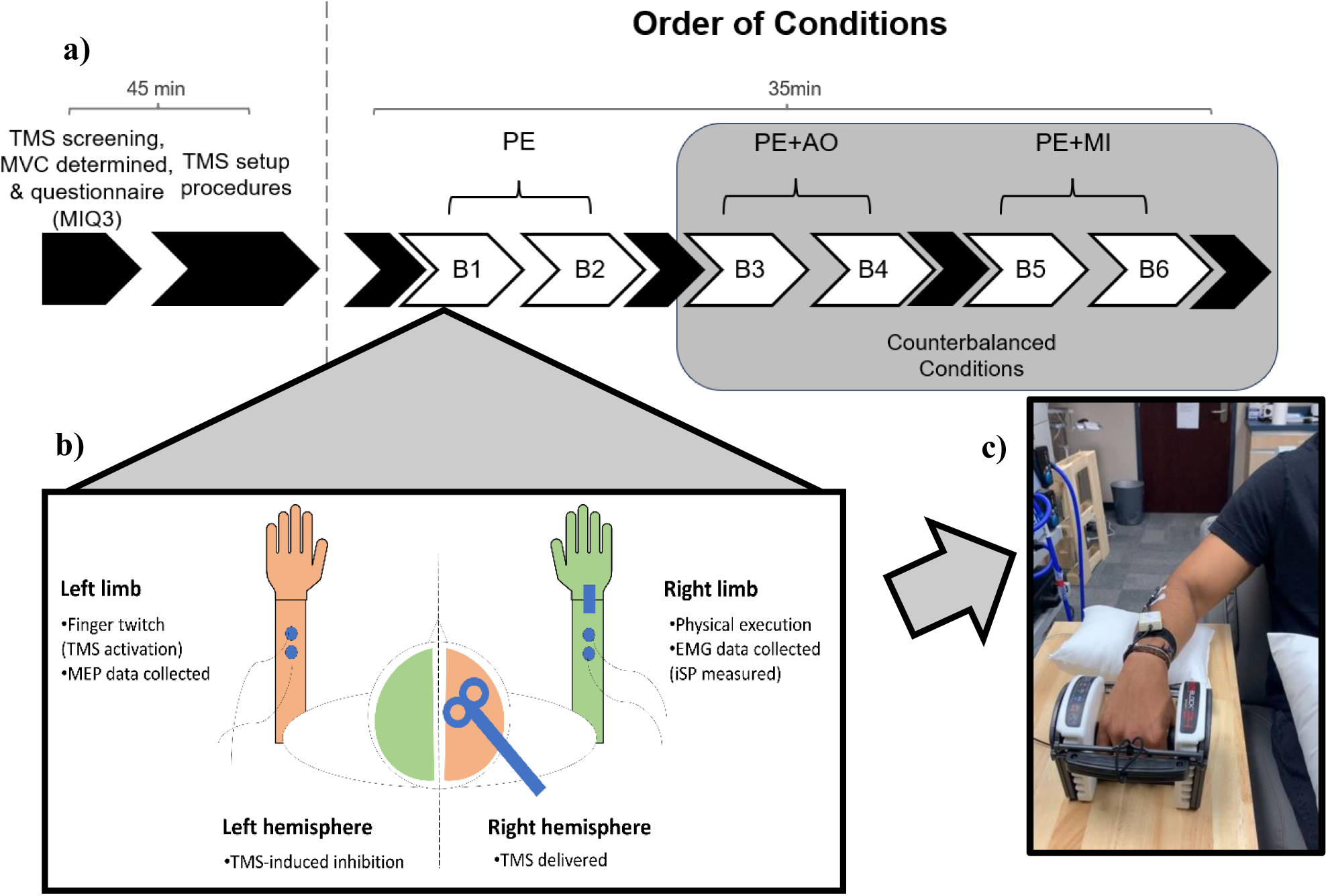
A) Timeline of the study session. Following background and setup procedures, participants performed two blocks of the PE alone condition. The PE alone condition was followed by two blocks of the concurrent action conditions (PE+AO or PE+MI), which were delivered in a counterbalanced order across participants. B) Experimental setup during each block of the study session for a right-handed participant. Single-pulse TMS was delivered and colour matching indicates contralateral hemisphere control of each limb (i.e., left hemisphere controlling right limb contractions). Outputs of each limb (collected through EMG) are also illustrated. C) One frame from the video (i.e., visual stimuli) presented from an external (3^rd^ person) perspective during the PE+AO condition. The electrode placement over the dominant forearm (extensor carpi radialis) can also be seen in this video, as well as the hand weight used in the physical task.

Alterations in IHI ultimately reflect changes in the activity of GABAergic circuitry which mediate connections between the primary motor cortices. When IHI is suppressed (indexed through a decreased iSP duration), this is thought to reflect ‘interhemispheric cooperation’ and/or an attenuation of a contralateral pattern of motor control.

Here, we aimed to assess IHI during covert action states, through a novel method of concurrently combining an active unilateral contraction with either MI or AO. We used this method to address our overarching objective of determining whether IHI is modulated during combined overt and covert action states, compared to when physical execution is performed alone, and to also further our understanding of the underlying neurophysiological processes occurring during the ‘execution’ of covert action states. Participants performed a force production task involving concentric wrist extensions with their dominant hand; either alone (i.e., physical execution/PE) or concurrently with either motor imagery (PE+MI) or action observation (PE+AO). In line with neuroimaging work suggestive of different patterns of brain activation for covert in comparison to overt action states, we predicted that IHI would be suppressed, indicated by decreased iSP duration, when these covert forms of practice were performed concurrently with physical execution. This suppression would indicate increased interference between the left and right motor cortices when additionally imagining the performed action or additionally seeing the action being performed by someone else.

There is also evidence to suggest that MI and AO will have different additive effects when combined with physical execution. MI is thought to be more similar to physical execution than AO, potentially having less of a suppression effect than AO (Hardwick *et al*., 2018; Kim et al., 2017; Sakamoto et al., 2009; Vogt et al., 2013; Kraeutner et al., 2023). Further, it has also been suggested that MI has different execution demands than physical execution, which might show up in an increased need for cross-hemisphere cooperation (Glover & Baran, 2017). Taken together, we hypothesized that IHI would be more greatly supressed during concurrent AO versus concurrent MI. Alternatively, an absence of change in IHI from physical execution alone to concurrent action conditions, may instead suggest that the neural representation governing control of these action states is functionally equivalent across the planning and execution of these actions, particularly at the level of cross-hemisphere projections that ‘quiet’ the ipsilateral hemisphere.

## METHODS

### Participants

Twenty-five typical adults were recruited for the study. Sample size was determined via previous work and a power analysis conducted using free statistical software (G*Power 3.1). We initially used a large effect size (ηp^2^ = .23), calculated from previous work evaluating interhemispheric inhibition in healthy individuals using similar TMS protocols (Kuo et al., 2017; McGregor et al., 2013; Strauss et al., 2019). It was determined that 12 participants would achieve statistical power (using an rmANOVA; α =.05, β = .05, η^p2^ = .23, Critical F = 3.49). To account for the novelty of our design (i.e., the use of non-invasive brain stimulation to assess interhemispheric inhibition during concurrent action modalities using a within-design, with potential attrition related to repeated measures) and given that samples sizes reported in previous work ranged from 10-21 in healthy participants with issues related to power and/or sample size noted (for a review see Hupfeld *et al*., 2020), we increased our initial sample size to 25.

Participants were included if they understood English and had normal or corrected-to-normal vision. Participants were excluded for contraindications to TMS (determined via an established TMS screening form; Rossi et al., 2009) and for having any neurological diagnosis or motor impairment that would impact fine upper limb motor coordination. Participants were recruited through the institution’s Psychology Undergraduate Study Pool and broader community. Two participants were excluded for technical issues during their study sessions, leaving twenty-three participants in the final analyses (20 females, 3 males, 21 right-hand dominant participants (determined via self report), 2 left-dominant participants, and an average age of 21 years). All participants provided written informed consent before participating in the study.

Before completing the behavioural task (below), participants’ motor imagery ability was characterized via the Movement Imagery Questionnaire-3^rd^ Version (MIQ-3; Williams et al., 2012). The MIQ-3 consists of twelve movements which are each first physically performed and then are imagined being performed based on three different criteria; i) internal-visual (which is imagining yourself perform the action from the 1^st^ person perspective as if ‘through one’s own eyes’), ii) external-visual (which is imagining yourself perform the action from the 3^rd^ person perspective, as if you were watching yourself on a video) and iii) kinesthetic (which is imagining the feelings and sensations associated with doing the action). Following their imagined performance, participants rated their imagery on a scale from 1 to 7 for how well they were able to see (visual dimensions) or feel (kinesthetic dimension) themselves performing the movement. Participants had a mean rating of 5.7 (*SD* = 1.3) on the internal visual imagery scale, 5.5 (*SD* = 1.3) on the external visual imagery scale, and 5.9 (*SD* = 1.0) on the kinesthetic imagery scale.

### Behavioural task

The behavioural task involved a concentric wrist extension with the dominant hand, performed with an adjustable PowerBlock (PowerBlock USA) weight. The weight was adjusted to be ∼50% of each participant’s maximum voluntary contraction (MVC). To determine MVC, each participant performed single repetitions of the concentric wrist extension task starting with the 3lb adjustable weight. After each successful repetition (full extension), an additional 3lbs were added until the participant could no longer perform a successful repetition. In this instance the weight of their last successful repetition was considered their MVC.

Participants performed the task under three conditions: physical execution alone (PE), physical execution and concurrent motor imagery (PE+MI), and physical execution and concurrent action observation (PE+AO). Participants performed the task with their arm 90 at degrees of abduction with the elbow at 90 degrees of flexion (i.e., beside their body such that their arm was in their periphery), beginning from a position where their arm was resting comfortably beside them on a padded (pillowed) arm rest (Figure 1). During the PE condition, participants were cued to perform the task from a custom developed program in Presentation (Neurobehavioral Systems Inc, USA) on a screen positioned in front of each participant. Each trial lasted 13 sec, comprised of a 4-second contraction and 9 seconds of rest. Visual and auditory cues were used to provide participants a 3 second countdown, indicating when participants should begin and hold the contraction and then rest (returning the weight to the start position). During the PE+MI condition, participants were additionally instructed to perform kinesthetic MI during the contraction, adopting a 1^st^ person perspective and focusing on imagining the sensations associated with performing the movement. Participants were instructed to perform MI with their eyes closed. During the PE+AO condition, participants additionally observed a video of the action played from an external visual, 3^rd^ person perspective. The video was of a person holding the same adjustable weight set to 9lbs, who also contracted their right arm (as seen in Figure 1c), which played simultaneously whilst participants physically performed the action.

Participants performed two consecutive blocks of each condition comprised of eight trials per block, with all participants beginning with two blocks of PE. The order that the PE+MI and PE+AO conditions were given were then counterbalanced across participants (see Figure 1). At the end of each block, ratings of perceived exertion (RPE; Williams, 2017) were obtained, rated on a scale from 6-20 (6 = no physical exertion and 20 = maximum physical exertion). RPEs were collected as a control measure to examine possible fatigue associated with repeated contractions.

### Transcranial Magnetic Stimulation (TMS)

TMS is a well-established and reliable technique which can be used to index corticospinal excitability (Hallett et al., 2000). Here we used single-pulse TMS to index interhemispheric inhibition during the three action conditions as described above. A neuronavigation system (Brainsight2^TM^, Rouge Research Inc., Montreal, Canada) was used to guide the positioning and orientation of the coil over the primary motor cortex. Each participant sat in a chair with their dominant hand beside them resting on a table (with a pillow) and their non-dominant hand resting on their lap. Electromyography (EMG; 1401 and Power 1902 Amplifier, Cambridge Electronics Design, UK) was recorded from electrodes attached to the extensor carpi radialis (ECR) muscle of each limb to allow measurement of both the iSP (in the ipsilateral limb to the pulse) and motor evoked potential (MEP) elicited in the contralateral limb. EMG signals were extracted using the neuronavigation EMG module (Brainsight2^TM^ EMG Isolation Unit and Amplifier pod, Rouge Research Inc., Montreal, Canada).

Before the task began, the motor “hotspot” and resting motor threshold (RMT; minimum stimulator output needed to elicit 5/10 MEPs ≥ 50 uV) was determined by following previously established protocols for determining the hotspot location in each participant with TMS (Kleim et al., 2007). A premade 5 by 5 grid (25 total points) was placed over the region of the brain where the primary motor cortex is located using the neuronavigation system. Starting in the centre of the grid at point (2, 2) and moving outward, the TMS was set to an initial stimulator output of 60% to find the point of the grid that exhibited the greatest MEP obtained from the contralateral ECR. Once the hotspot location was determined, the TMS stimulator output systematically reduced to determine RMT.

During each trial of the behavioural task, single-pulse TMS was delivered over the ipsilateral hemisphere (e.g., right, for a right-handed individual) during the 4sec contraction period at a fixed 2.5sec following the cue at 130% of RMT. To index interhemispheric inhibition, we measured the ipsilateral silent period (iSP; latency period between the onset and offset of any inhibitory signals being sent from one side of the brain to another during the active contraction). MEPs obtained during each trial (from the contralateral, resting limb), were used to control for potential changes in corticospinal excitability observed across the session.

### Data analysis

#### Interhemispheric inhibition

The ipsilateral silent period (iSP) was our primary outcome measure, which was quantified using established procedures (Garvey *et. al.*, 2001; Hupfeld *et al.,* 2020). In brief, using a customized Matlab script, EMG traces were first rectified. Each trial was quality-checked by a single rater (KV), with trials removed for excessive noise in the EMG signal or an inconsistent EMG signal (i.e., erroneous movements made by the participant). The rectified EMG traces were then averaged across each block (Hupfeld et al., 2020). The average EMG amplitude in the 100 ms period preceding the TMS pulse was noted as the mean pre-stimulus level with a threshold determined via Mean Consecutive Difference approach used to identify the iSP (Hupfeld et al., 2020). Onset of the iSP was defined as the point at which the post-stimulus (i.e., after the TMS pulse) EMG fell below the threshold for five consecutive data points. Offset of the iSP was defined as the point at which the post-stimulus EMG returned above the threshold for five consecutive data points. Duration of the iSP was then calculated for each trial as iSP offset minus iSP onset.

A linear mixed effects (LME) model was conducted to assess differences in iSP across the conditions. We included both Condition (3 levels pertaining to the three conditions) and condition Order (2 levels; PE+AO first vs. PE+MI first) as fixed effects, and participant as a random effect. Preplanned orthogonal Helmert contrasts were applied to the factor Condition to test whether iSPs differed between (i) control (PE) and experimental conditions (PE+MI and PE+AO) and (ii) between the two experimental conditions (PE+MI vs PE+AO).

#### Corticospinal excitability

The peak-to-peak amplitude of MEPs evoked during each trial (from the contralateral, resting limb), were used to index corticospinal excitability. The EMG trace for each trial was quality-checked by a single rater (KV), with trials removed for excessive noise or indications that the resting limb had moved. MEPs remaining in final analyses were then normalized to the PE alone condition (control; concurrent minus PE).

An LME model was again applied to test whether normalized MEPs differed between the two experimental conditions. Condition (2 levels, PE+MI vs PE+AO) and condition Order (2 levels; PE+AO first vs. PE+MI first) were entered as fixed effects and participant was entered as a random effect.

#### Perceived exertion

Ratings of Perceived Exertion (RPEs) were recorded throughout the session to assess potential fatigue induced by repetitive contractions. A third linear mixed effects model was conducted to assess differences in RPE across the three conditions, as used for the iSP analysis.

## RESULTS

The average weight (50% MVC) used in our study was 6lbs. Specifically, participants who engaged in AO first (i.e., PE+AO *preceding* PE+MI; n = 11, aged 22 ± 3.6, 10 female, 11 right-handed) had an average 50% MVC of 5.5 ± 1.2. Participants who engaged in MI first (i.e., PE+AO *after* PE+MI; n = 12, aged 21 ± 1.7, 10 female, 10 right-handed) had an average 50% MVC of 6.8 ± 1.9.

### Interhemispheric inhibition

Across all participants, four trials were excluded from the PE+MI condition (1.1% of total trials for that condition), and three trials were excluded from the PE+AO condition (0.82% of total trials for that condition). No trials were excluded for the PE alone condition.

Figure 2 illustrates mean iSP duration across the two blocks for participants who did AO first and for participants who did MI first. Based on the LME model with pre-planned contrasts, there was a significant difference between the two experimental conditions only (*p =* .005; see Table 1). No difference was observed when the PE control condition was compared to the mean of both experimental conditions (*p* = .13). Generally, the difference between experimental conditions resulted in a reduction in iSP duration in AO versus MI, but as can be seen in Figure 2, this general effect was dependent on the order of conditions. Only when the PE+AO condition was first, did the AO condition show a shorter iSP duration compared to MI (PE+MI *minus* PE+AO; *d* = 0.67), which was not present when the MI condition was first (PE+MI *minus* PE+AO; *d* = −0.29). This order effect was statistically evidenced by an interaction of Condition and condition Order (*p* = .014).

**Figure 2.**
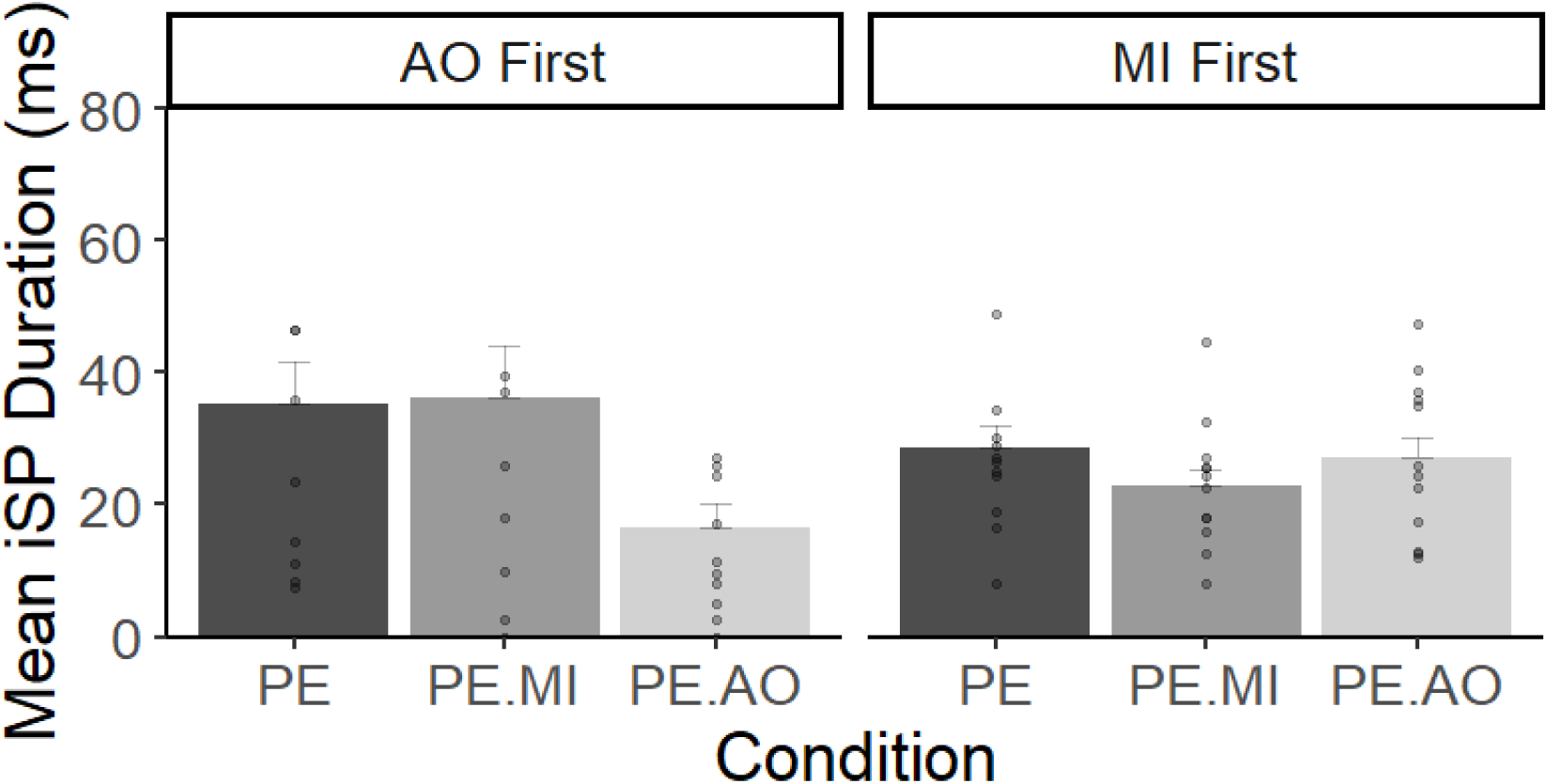
Mean iSP duration across both blocks of each of the three conditions, for participants who did PE+AO prior to PE+MI (left) or PE+MI prior to PE+AO (right). All participants performed the PE condition first. Errors bars represent standard error of the mean for each condition.

**Table 1:**
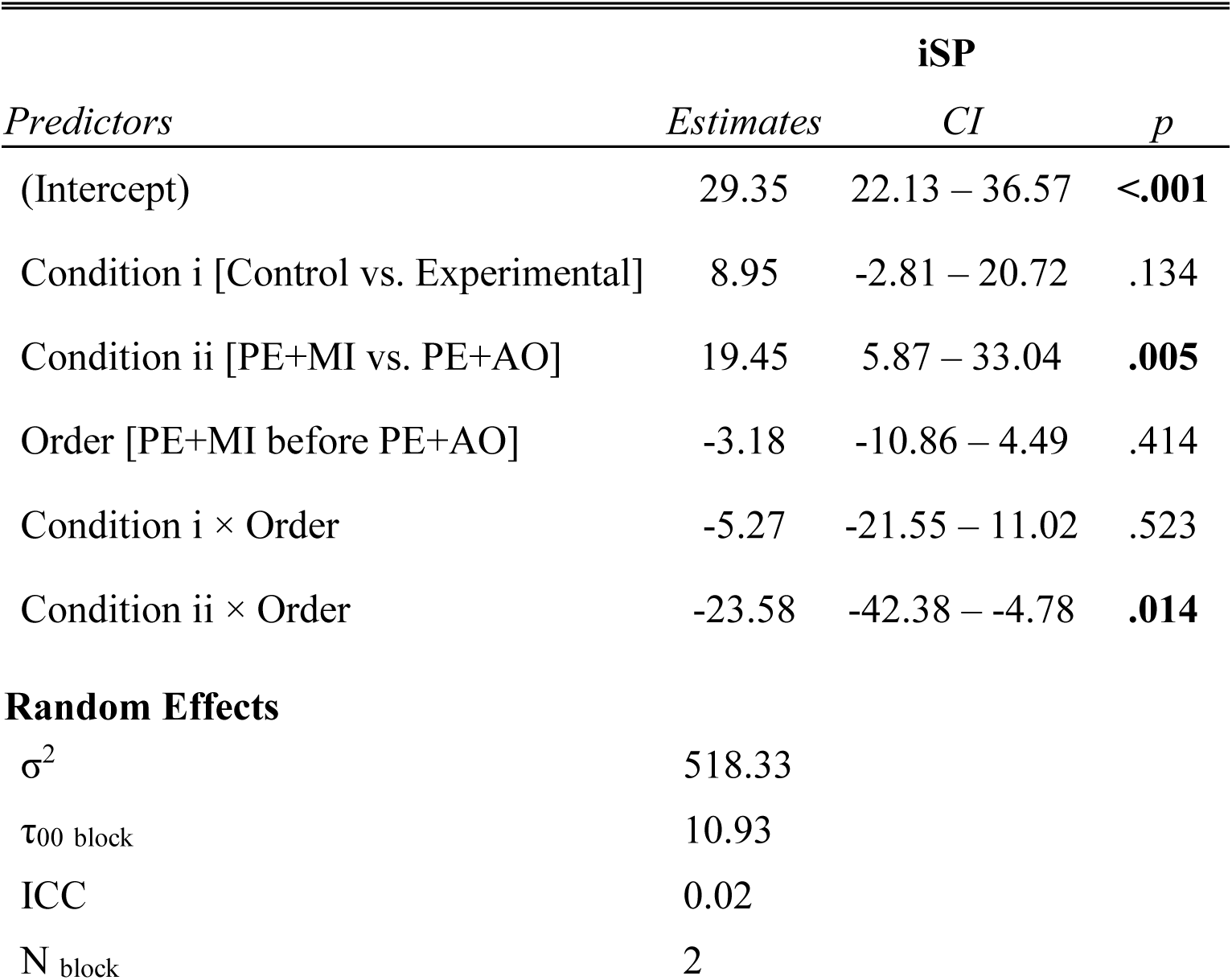
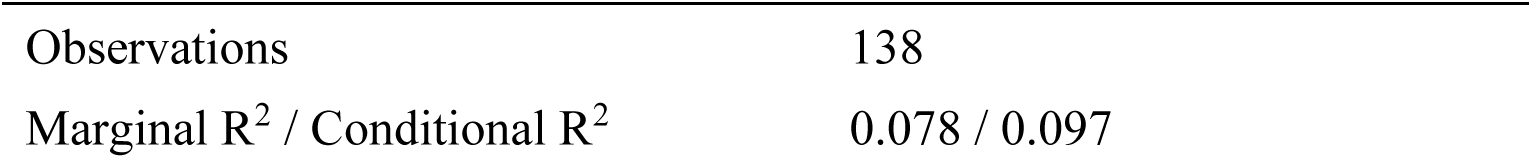
Linear mixed effects model conducted on iSP duration as a function of condition and order.

| Predictors | iSP |  |  |
| --- | --- | --- | --- |
|  | Estimates | CI | p |
| (Intercept) | 29.35 | 22.13 – 36.57 | <.001 |
| Condition i [Control vs. Experimental] | 8.95 | -2.81 – 20.72 | .134 |
| Condition ii [PE+MI vs. PE+AO] | 19.45 | 5.87 – 33.04 | .005 |
| Order [PE+MI before PE+AO] | -3.18 | -10.86 – 4.49 | .414 |
| Condition i × Order | -5.27 | -21.55 – 11.02 | .523 |
| Condition ii × Order | -23.58 | -42.38 – -4.78 | .014 |
| **Random Effects** |  |  |  |
| $\sigma^2$ | 518.33 | | |
| $\tau_{00}$ block | 10.93 | | |
| ICC | 0.02 |  |  |
| N<sub>block</sub> | 2 |  |  |

|  |  |
| --- | --- |
| Observations | 138 |
| Marginal $R^2$ / Conditional $R^2$ | 0.078 / 0.097 |

### Corticospinal excitability

Four participants were removed from MEP analyses, due to EMG activity that preceded the MEP (indicating overt movement in the resting limb) and/or excessive noise in the signal. Across all remaining participants (n=19), one trial was excluded from the PE+AO condition, which accounted for 0.27% of total trials for that condition. No trials were excluded from the other two conditions.

Figure 3 illustrates normalized MEP amplitude (normalized to the PE alone condition). No difference in normalized MEP amplitude was observed between the two experimental conditions based on the LME model (*p =* .62; Table 2). There was also no effect of condition Order (*p*= .56) nor an interaction between the two (*p* =.68).

**Figure 3.**
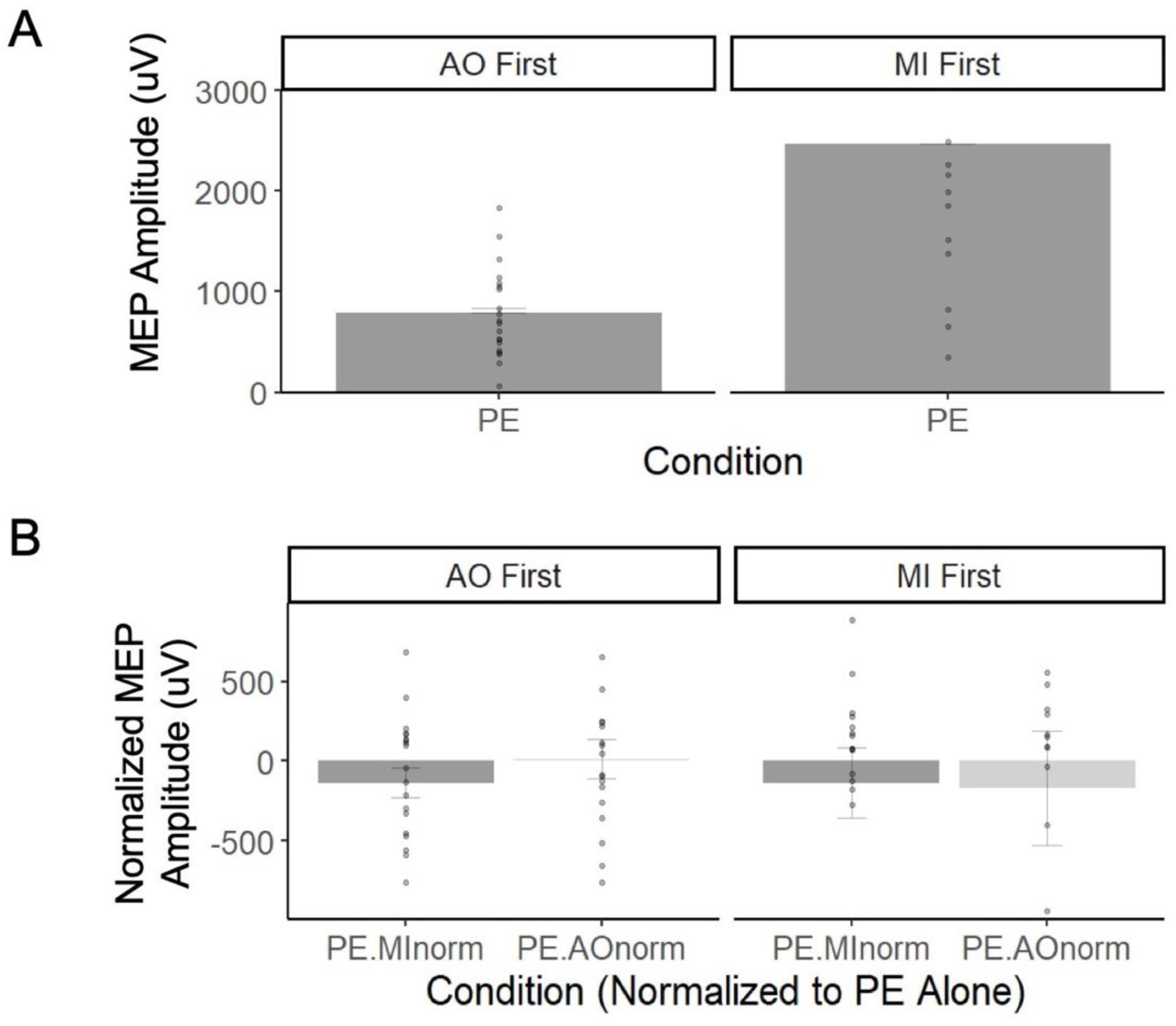
MEP data subdivided based on participants that did PE+AO first (left) or PE+MI first (right). A) Mean MEP amplitude for the PE condition. B) Mean normalized MEP amplitude for the two experimental conditions (with MEP amplitudes for each experimental condition normalized to the PE alone condition for each participant). Individual participants are represented by the data points, and errors bars represent standard error of the mean for each condition.

**Table 2:** Linear mixed effects model conducted on normalized MEPs as a function of condition and order.

| *Predictors* | **Normalized MEPs** |  |  |
| --- | --- | --- | --- |
|  | *Estimates* | *CI* | *p* |
| (Intercept) | 11.48 | -409.07 – 432.03 | .957 |
| Condition [PE+MI] | -149.94 | -744.69 – 444.80 | .617 |
| Order [PE+MI before PE+AO] | -180.67 | -791.71 – 430.37 | .557 |
| Condition [PE+MI] ×<br>Order [PE+MI before PE+AO] | 181.95 | -682.19 – 1046.09 | .676 |
| **Random Effects** |  |  |  |
| $\sigma^2$ | 889240.98 | | |
| $\tau_{00 \text{ block}}$ | 0.00 | | |
| $N_{\text{block}}$ | 2 | | |
| Observations | 76 |  |  |
| Marginal $R^2$ / Conditional $R^2$ | 0.006 / NA | | |

**Table 3:**
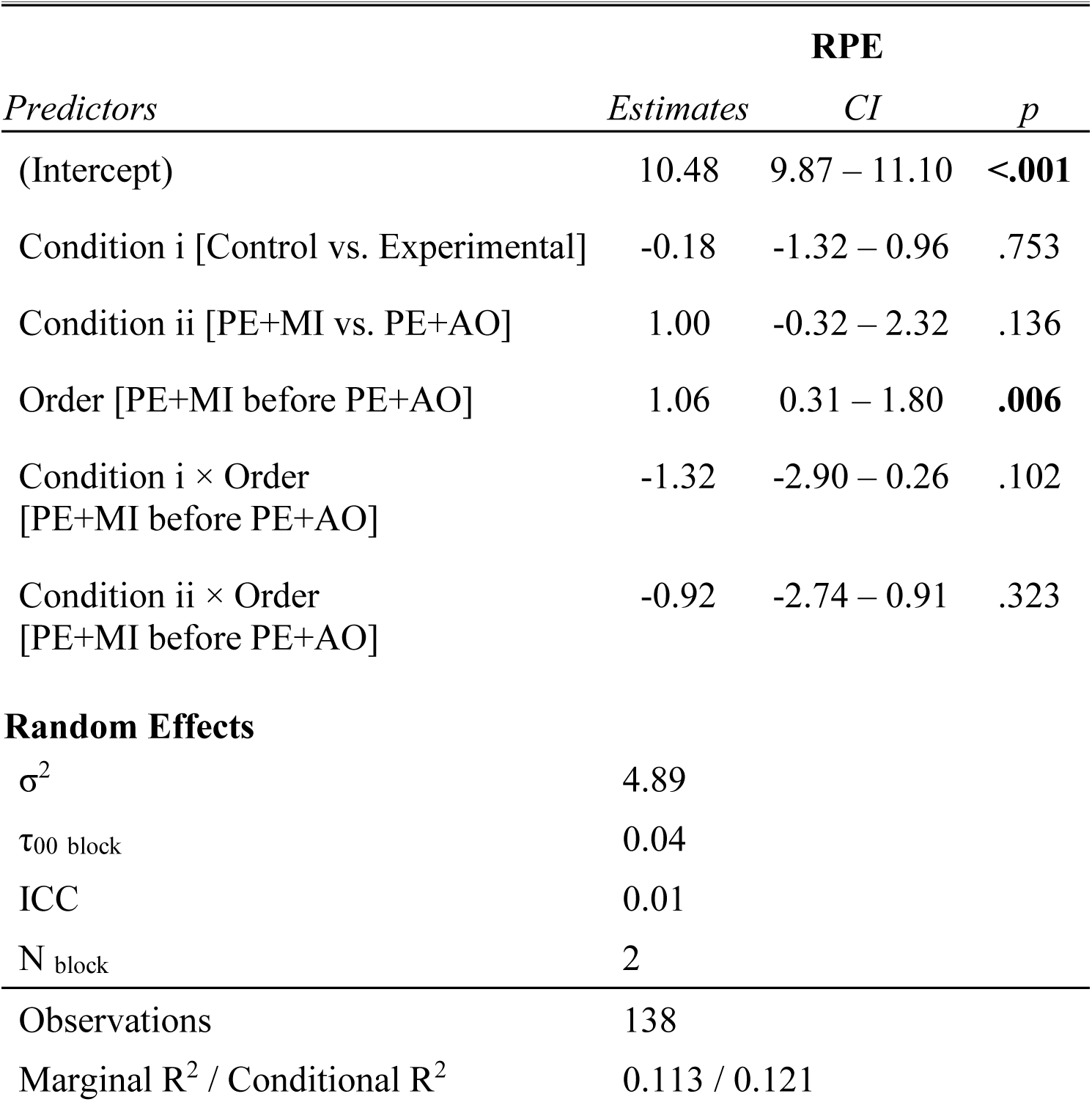
Linear mixed effects model conducted on RPEs as a function of condition and order.

### Perceived Exertion

All ratings were included in the analysis. Figure 4 illustrates RPE ratings by Condition and condition Order. There were no differences as a function of condition based on the pre-planned contrasts. There was an effect of condition Order (*p* = .006), due to higher ratings of RPE for participants who engaged in PE+MI before PE+AO.

**Figure 4.**
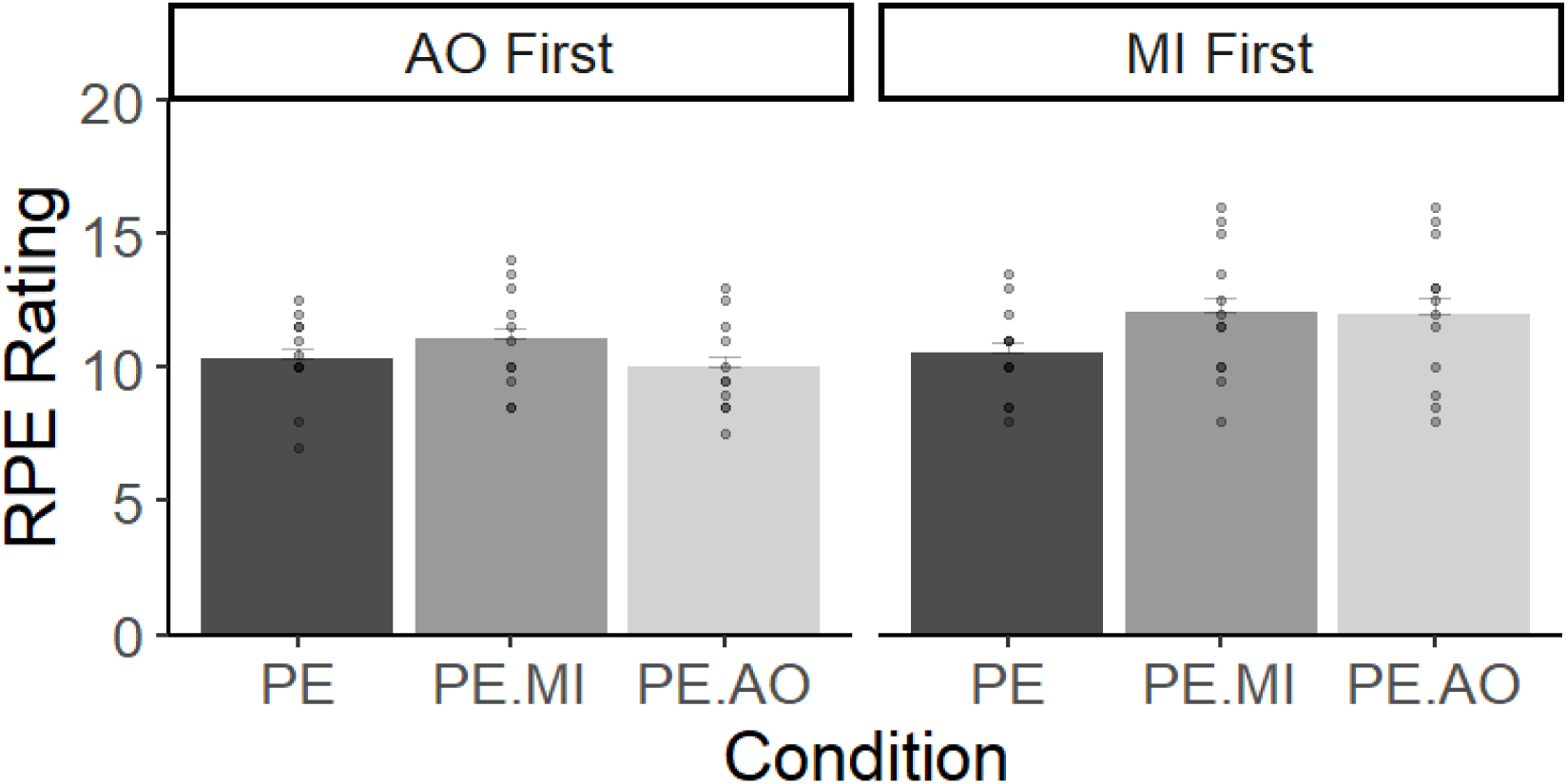
Mean RPE Ratings across both execution blocks for all three conditions, subdivided based on participants who did PE+AO first (left) and PE+MI first (right). Error bars represent standard error of the mean for each condition.

## DISCUSSION

In this study we used a within-participants’ design to probe modulations of IHI when physical execution of a single arm contraction was concurrently paired with action observation (AO) or motor imagery (MI). In partial support of our hypotheses, IHI was decreased when AO was paired with physical execution, compared to when physical execution was performed alone. However, this decrease in IHI was dependent on the order that conditions were conducted and was only evident when the AO condition was performed before the MI condition. Contrary to our hypotheses, MI did not modulate IHI. We discuss these IHI data in the context of our other measures as well as in comparison to prior neuroimaging and theoretical work speaking to the potential equivalence between covert action states of AO and MI and physical execution.

### Interhemispheric cooperation vs. inhibition

When engaged in concurrently with physical execution, AO led to a suppression in IHI (indexed by a decrease in iSP duration). This result suggests that AO requires bilateral recruitment of the motor cortices, relying on ‘interhemispheric cooperation’ rather than IHI, in comparison to physical execution and MI. Other researchers have shown bilateral activation of the motor cortices during AO during various unilateral tasks (e.g., Alhajiri et al., 2018; for meta-analytic work see Hardwick et al., 2018). For instance, Alhajiri et al. (2018) used electroencephalography (EEG) to assess mu rhythm suppression (thought to reflect modulation of sensorimotor areas underlying movement) during AO (vs. physical execution) of a joystick-tracing task. During AO, mu rhythm measured from electrodes localized over the contralateral hemisphere and midline was suppressed (similarly to suppression observed during physical execution of the task). Although suppressed to a lesser extent, mu rhythm recorded from the electrode localized over the ipsilateral hemisphere was also suppressed, indicating a low level of activation of the ipsilateral motor cortex during observation of a unilateral task.

In behavioural work, there is evidence that concurrent AO can interfere with physical execution, although in these tasks the AO is of an action different to the one being performed (e.g., Kilner et al., 2003). For example, when participants concurrently observed actions that were either incongruent or congruent with the action they were physically executing, movement variance increased during concurrent AO of incongruent actions (Kilner et al., 2003). Although we had no reason to suspect that concurrent AO in our study would interfere with physical execution (as participants observed a congruent action), there is the possibility that it did produce a degree of interference, acting more like a secondary “monitoring” task.

Motor imagery, in contrast to AO, did not modulate IHI when performed concurrently with physical execution. Two explanations exist for the lack of differences. First, it is challenging to monitor performance of MI given its concealed nature and the subjective nature of many tools used to assess MI ability (Boe & Kraeutner, 2017); thus, we are unable to confirm whether participants were able or actually performed MI during the contraction. It may be challenging to both imagine feeling the movement and perform a live contraction. Specifically, there is potential that participants found it difficult to imagine the kinesthetic properties of the task, due to concomitant actual kinesthetic feedback. In this case, participants may have focused (internally) on the kinesthetic aspects of the physical performance (Wulf, 2013). We did not ask people to rate their ability to imagine after each trial (in order to keep the timing between conditions consistent); which in hindsight, may have provided a better manipulation check concerning ability to concurrently imagine and act. If we look at the RPE data as an index of the effort required of the task, although there was no overall difference across conditions, the order that conditions were performed did matter. When participants performed MI before engaging in AO, they rated the MI condition as more effortful. The requirement to additionally image whilst contracting was seen as more ‘physically’ effortful when it was not preceded by any practice. Future work using an approach that perhaps tests or dissociates these imagined and actually experienced sensations (i.e., performing MI of a weight different to 50%MVC) may be needed to determine if MI can modulate IHI (rather than just an enhanced internal focus on attention onto the movement, which did not impact IHI here).

A second reason for the lack of difference between MI and physical execution may be that MI does not rely on the recruitment of the primary motor cortices (Hétu et al. 2013). Activation of the motor cortices during MI is not consistently observed across neuroimaging work (Hétu et al., 2013), thus, the inhibitory pattern of control maintained in our PE+MI condition could simply be a result of a lack of any effect of MI on motor cortical activity over and above that seen from physical execution. In line with the theory that MI relies more strongly on cognitive processes not evidenced during physical execution (Glover & Baran, 2017), past work suggests that motor cortical activity observed during MI is modulated by an indirect inhibitory pathway from posterior parietal regions (Lebon et al., 2012). Specifically, following a subthreshold conditioning stimulus applied to the posterior parietal cortex, Lebon et al., 2012 showed that resultant MEPs elicited from the contralateral motor cortex were suppressed relative to rest, indicating an inhibitory role of the parietal cortex (i.e., related to inhibiting overt movement) during MI. Our data supports this inference, in that MEPs obtained during concurrent action states were suppressed relative to PE alone (‘negative’ normalized MEPs).

It is then perhaps counterintuitive that we did not observe a difference in corticospinal excitability (indexed via assessing MEP amplitudes elicited from the ipsilateral hemisphere) between our experimental conditions, given past work showing that corticospinal excitability during AO is decreased relative to MI (Wright et al., 2018). While work directly comparing parietal-motor connectivity between MI and AO suggests this parietal-motor pathway is less relevant for AO (Feurra et al., 2011), others have shown that motor cortical activity observed during AO is caused through excitation of premotor areas that directly project to the motor cortex (Gallese et al., 1996; Loporto et al. 2011). Taken together with our findings, suggest that MI does not *rely* on the motor cortices, and it is likely that motor cortical activity observed during AO and MI is modulated through different cortical pathways.

### Order effects associated with motor imagery

Here, we observed an asymmetry in the extent to which IHI occurred during PE+AO, depending on the order of conditions. There was a decrease in IHI when AO was combined with physical execution, compared to being performed alone, but only when the PE+AO condition occurred first. This was an unexpected effect and hence our reasons for these order effects are only speculative. One potential explanation for the lack of any difference between the PE and the addition of concurrent AO when MI was performed first may be because MI was incidentally primed during the PE+AO condition. If that is the case, this would suggest that MI negates any influence of AO, changing the focus of attention away from the video image for example. However, equally plausible is that merely performing the AO condition last may lessen the focus on the concurrent video and as such prevent us seeing a moderation of IHI.

There has been considerable research interest directed to determining the combined effects of AO+MI for practice, with the assumption that these covert states activate different processes that together can augment performance and learning experiences for individuals (for reviews see Eaves *et al*., 2022; Scott *et al*., 2022). Our data are generally in alignment with conclusions from this work, showing that there are behavioural as well as neurophysiological control related differences between AO and MI. Interestingly, in some of this literature where AO and MI effects have been compared separately to those when they are combined, protocols have deliberately avoided introducing MI prior to AO to prevent spontaneous (i.e., priming effects of) MI during AO (Emerson et al., 2022; Scott et al., 2019; Scott et al., 2020). Thus, it is difficult to assess AO after MI as it appears that individuals continue to automatically engage in MI even when no explicit instructions are given to do so (e.g., Maslovat et al., 2013).

### Considerations

There were a few limitations of the current design that impact conclusions and prompt further study. The first is that it is possible that induced fatigue may have led to temporary reductions and disruption of functional connectivity between motor cortices of the hemispheres (Peltier et al., 2005). However, to mitigate fatigue-related confounds, the number of trials participants performed per block and across the study session was minimized, with rest provided between trials and blocks. Further, participants’ perceived exertion (assessed by RPEs) was consistent across conditions. However, RPEs were greater for participants who performed PE concurrently with MI before engaging in concurrent AO. This greater RPE was therefore an effect related to the order of conditions, with those who performed MI earlier on rating the overall session as being more “effortful”. These RPE data may indicate that MI itself induced additional fatigue or a propensity for participants to conflate task difficulty (from combining MI with physical execution) with increased perceived exertion and further suggesting that these individuals continued to engage in MI during the PE+AO condition.

A potential limitation of the current design was the focus on a single viewing perspective during the concurrent AO condition. Participants only viewed videos shown from a third-person perspective. In this way, the observed action was performed with the same limb as the participant, but it would appear as being on the opposite side (i.e., when facing a person performing an action with the same limb). For right-handed individuals, the delivery of AO in a similar perspective as in our study has been shown to elicit greater MEPs in the right as compared to the left limb (Sartori et al., 2013, 2014) and was the reason we chose to show the video from this perspective. However, it will be necessary in future work to test different perspectives that are either anatomically or spatially compatible to the actions being produced by the participant to determine whether interhemispheric cooperation is a result of perspective, or a general effect associated with concurrent AO. Moreover, while the majority of participants in our study identified as right-handed, two were left-handed and there is evidence that left-handed individuals show an innate tendency to represent actions in the left limb regardless of the observed effector (Sartori et al., 2013, 2014). There was, however, no evidence in our study that the left-handed individuals responded differently to the right-handed, in terms of condition and condition order effects.

In conclusion, here we investigated the potential modulation of IHI through concurrent performance of covert actions (MI, AO) with overt movement. Our data suggest a separation between AO and MI, as concurrent PE and AO led to suppressed IHI (i.e., enhanced interhemispheric cooperation), yet IHI was not modulated when MI was performed concurrently with PE. These data provide further evidence to suggest additional recruitment of the ipsilateral motor cortex during AO, and in-turn may help to explain interference effects produced by concurrent AO during online motor control. However, these effects were dependent on the order of conditions and only when the AO condition was performed first (rather than following MI), was evidence of modulation of IHI observed. One reason for these order effects may be related to the overriding influence of MI that continues to impact AO in the absence of any explicit instructions to engage in MI. In future work, it will be important to consider the effects of different viewing perspectives adopted during AO when assessing underlying neurophysiology and employing other measures to assess the ability to engage in imagery during each trial. Overall, this study furthers our understanding of the neurophysiological processes that are associated with AO and MI, when performed concurrently with PE and shed light on different control processes underpinning AO and MI and their assumed functional equivalence.

## ACKNOWLEDGMENTS

This work was supported by funding awarded to SK through the Natural Sciences and Engineering Research Council (NSERC; Discovery Grant). KV was supported by an Undergraduate Student Research Award (NSERC USRA). CP is supported by a Postgraduate Scholarship (NSERC PGS D).

## AUTHOR CONTRIBTUIONS

KV: data curation, methodology, formal analysis, visualization, writing – original draft, writing – review & editing.

MS: conceptualization, methodology, formal analysis, visualization, writing – original draft, writing – review & editing.

CP: methodology, formal analysis, writing – review & editing. KS: data curation, methodology, writing – review & editing.

NH: conceptualization, methodology, writing – review & editing.

SK: conceptualization, methodology, writing – original draft, writing – review & editing, funding acquisition, supervision, project administration.

## Notes

### Competing Interest Statement

The authors have declared no competing interest.

